# Dose-dependent activation of Syk and SHIP1 by LynA and LynB at steady state creates a threshold for macrophage signaling in the absence of receptor engagement

**DOI:** 10.64898/2026.02.25.708040

**Authors:** S. Erandika Senevirathne, Carter E. Sellner, Silvia Toledo Ramos, Tanya S. Freedman

**Affiliations:** Graduate Program in Molecular Pharmacology and Therapeutics, University of Minnesota, Minneapolis, MN 55455, United States; Department of Pharmacology, University of Minnesota, Minneapolis, MN 55455, United States; Center for Immunology, University of Minnesota, Minneapolis, MN 55455, United States; Masonic Cancer Center, University of Minnesota, Minneapolis, MN 55455, United States

**Keywords:** Src-family kinase, Inositol phosphatase, Syk kinase, Steady-state regulation, Macrophage

## Abstract

LynA-knockout and LynB-knockout mice, each expressing only one of two isoforms of the Src-family kinase (SFK) Lyn, have differential progression to autoimmunity. It is unclear, however, whether isoform-specificity or Lyn dose underlies differential signaling in the single-isoform knockouts. To address this question, we generated a series of Lyn-knockout mice with a varying LynA and LynB expression and tested macrophage signaling in response to pharmacological pan-SFK activation. We found that the magnitude of initiating signaling is a function of the combined basal expression of LynA and LynB, with the two isoforms equally capable of phosphorylating positive-regulatory Syk and negative-regulatory SHIP1. While expression of either isoform restored basal and SFK-initiated downstream signaling, WT-like levels of Erk and Akt signaling were enabled by expression of any amount of Lyn and insensitive to further upregulation of either isoform. Thus, either LynA or LynB expression at steady state leads to balanced activation of positive- and negative-regulatory signaling, setting a maximal response in the absence of a true microbial encounter.

**Summary Sentence:** Total expression of LynA and LynB determines the steady-state phosphorylation of the activating kinase Syk and the inhibitory phosphatase SHIP1, capping signaling in the absence of a microbial encounter.

## Introduction

Src-family kinases (SFKs) regulate signaling responses in many receptor pathways, including immunoreceptor tyrosine-based activation motifs (ITAMs) and immunoreceptor tyrosine-based inhibitory motifs (ITIMs)^1^. Phosphorylation of ITAM tyrosine residues by activated SFKs leads to recruitment and partial activation of the down-stream kinase Syk^1-3^, where SFK-mediated phosphorylation of tyrosine (Y) 352 in the SH2-kinase interdomain-B linker of Syk facilitates full activation of Syk by activation-loop autophosphorylation^4^. Together, activated SFKs and Syk phosphorylate actin-remodeling proteins, adaptors, and second-messenger-generating enzymes, which ultimately lead to Erk and Akt pathway activation and a broad antimicrobial response^2,3,5^. Conversely, phosphorylation of ITIMs by SFKs leads to the recruitment of phosphatases, such as the inositol phosphatase SHIP1^1^, which limits immune cell responses by dephosphorylating PIP_3_^6^ suppressing further signaling.

The SFK Lyn initiates signaling through both ITAM and ITIM pathways^7^. Global^8-10^, B-cell-specific^11^, and myeloid-cell-specific^12,13^ Lyn-knockout (KO) mice and Lyn-overexpressing mice^14^ develop autoimmune disease, highlighting the intricate balance of pro-inflammatory and suppressive functions of Lyn. Previous work from our group demonstrated that Lyn, but not the SFKs Hck and Fgr, suppresses basal activation of the Erk and Akt pathways in macrophages by phosphorylating SHIP1 and preventing accumulation of PIP_3_^15^—basal activation of SHIP1 was revealed as the key barrier between activation of SFKs and Syk at the membrane and initiating downstream antimicrobial signaling in the absence of an encounter with a receptor-binding particle. Interestingly, this basal rheostat function, not the degree of further phospho (p)SHIP1 induction, determined the strength of subsequent signaling^15^.

Lyn is expressed as two splice variants, LynA and LynB, which are produced by alternative splicing of the same gene, differing in a 21-amino-acid insert within the unique region of LynA that is lacking in LynB^16^. We previously reported that phosphorylation of Y32 in the unique-region insert of LynA, either by LynB or LynA itself, flags LynA for polyubiquitination by c-Cbl^17^ and subsequent activation-induced degradation^18^. Despite an early lentiviral-transduction study of LynA and LynB in to Lyn^KO^ in mast cells that suggested segregation of activating and inhibitory signaling functions to LynA and LynB, respectively^19^, and a unique role for LynA in aggressive cancers^20^, little is known about the functions of the two Lyn isoforms.

To test the independent contributions of LynA and LynB to immune regulation, our group used CRISPR/Cas9 gene editing to generate single-isoform LynA^KO^ and LynB^KO^ mice and found that LynB was the dominant isoform protecting against autoimmune disease^10^. Nevertheless, we found sexual dimorphism primarily in LynA^KO^ mice, and the cellular phenotypes for the single-isoform knockouts (used as F1 crosses with complete Lyn^KO^ to ensure WT-level expression of the remaining isoform) were distinct from each other and from complete Lyn^KO10^. A complicating factor in interpreting these results, however, is that, perhaps due to differences in c-Cbl expression^17^, the relative expression levels of LynA and LynB are subtly but reproducibly cell-type-specific^10^. Therefore, it was unclear whether the more severe phenotype in LynB^KO^ was attributable to loss of isoform-specific functions or to loss of the isoform more highly expressed in disease-promoting cells.

For this study we modulated LynA and LynB expression individually to reveal the relative contributions of isoform-specific functions and dose-dependent functions to macrophage signaling. To examine the intrinsic potential to phosphorylate substrates without the confounding complexities introduced by receptor engagement^12,15^, we used bone-marrow-derived macrophages (BMDMs) from transgenic mice lacking the endogenous SFK-inhibitory kinase Csk and expressing an analog-sensitive (AS) variant of the SFK-inhibitory kinase Csk, which can be selectively inhibited by the small-molecule inhibitor 3-IB-PP1, leading to robust, receptor-independent SFK activation^17,18,21,22^. We have previously used this approach to probe LynA-specific post-translational modification^17,18^, requirements for pro-inflammatory signaling^18^, and the basal requirement for Lyn (A + B) expression and SHIP1 phosphorylation in regulating macrophage signaling^15^. We now build on this framework to demonstrate that LynA and LynB both have the potential to activate Syk and downstream signaling. Most interestingly, we find that the combined expression level of LynA and LynB at steady state defines the setpoint of basal phosphorylation for SHIP1. This Lyn/SHIP1 steady-state rheostat sets an upper limit on downstream signaling and balances activation of the Erk and Akt pathways in the absence of receptor engagement.

## Methods

### Mice

All mice were derived on a C57BL/6 background. Homozygous LynA or LynB knockouts with supraphysiological expression of the remaining isoform were generated by breeding CRISPR-generated LynA-knockout mice or LynB-knockout (LynB^+/+^LynA^KO^ or LynA^+/+^LynB^KO^: UP)^10^. Hemizygous LynA or LynB knockouts with WT-like expression of the remaining isoform were generated by crossing homozygous LynA-knockout or LynB-knockout mice to neomycin (neo)-disrupted total Lyn^KO^ mice^23^, yielding animals that express the remaining Lyn isoform from a single allele (LynB^+/-^LynA^KO^ or LynA^+/-^LynB^KO^: PHYS)^10^. Lyn-knockout Csk^AS^ mice were generated by crossing Lyn^KO^, LynA^KO^, or LynB^KO^ to Csk^KO^Csk^AS^ transgenic mice^10,21^. Animal studies were performed in compliance with the University of Minnesota/American Association for Accreditation of Laboratory Animal Care and National Institutes of Health policy, under Animal Welfare Assurance number A3456-01 and Institutional Animal Care and Use Committee protocol 2506-43046A. All animals were housed in specific-pathogen-free conditions and supervised by licensed veterinarians.

### Preparation of BMDMs

BMDMs were prepared as previously described^18^. Bone marrow flushed from the femora and tibiae of 10-15-week-old male and female mice, subjected to hypotonic erythrocyte lysis, and cultured in untreated plates (Corning, Manassas, VA, USA) in Dulbecco’s Modified Eagle Medium (DMEM, Corning) supplemented with 10% heat-inactivated fetal bovine serum (FBS, Omega Scientific, Tarzana, CA, USA), 0.11 mg/ml sodium pyruvate (Corning), 2 mM penicillin/streptomycin/L-glutamine (Sigma-Aldrich, St. Louis, MO, USA), and 10% CMG-14-12-cell-conditioned medium as a source of macrophage colony-stimulating factor (M-CSF). After 7 days of culture, BMDMs were detached from plates with 5 mM ethylenediaminetetraacetic acid and replated at 1×10^6^ cells/well in untreated 6-well plates (Corning) in M-CSF-free medium supplemented with 25 U/ml IFN-γ (Peprotech, Rocky Hill, NJ, USA) for inflammatory priming overnight.

### Cell stimulation

BMDMs were treated with 10 µM 3-IB-PP1^15,17,18^ and incubated at 37°C before quenching on ice and lysing in sodium dodecyl sulfate (SDS) buffer (10% glycerol, 4% SDS, 50 mM dithiothreitol, and 128 mM Tris base, pH 6.8). After scraping cells and incubating 5 min at 37°C, lysates were collected and sonicated on a Bioruptor (Diagenode Inc, Denville, NJ, USA) at 50% duty for 3 min to shear DNA. Lysates were then boiled 15 min and stored at -20°C.

### Immunoblotting

Samples were separated on a 7% NuPage Tris-Acetate gel (Invitrogen, Carlsbad, CA, USA) and then transferred to Immobilon-FL PVDF membrane (EMD Millipore, Burlington, MA, USA). Revert Total Protein stain (LI-COR Biosciences, Lincoln, NE, USA) was used to quantify the total protein content in each lane. After destaining, membranes were cut as appropriate and blocked 1 h with TBS Intercept Blocking Buffer (LI-COR). Membranes were then incubated overnight at 4°C with primary antibodies, treated 1.5 h with secondary antibodies, and visualized on an Odyssey CLx near-infrared imager (LI-COR). Densitometry quantification was performed in ImageStudio (LI-COR), as described previously^24^.Antibodies were obtained from Cell Signaling Technologies (cs, Danvers, MA, USA), Abcam (ab, Cambridge, UK), or LI-COR (lc) as follows: rabbit anti-pSyk^Y352^ 65E4 cs2717, rabbit anti-pSHIP1^Y1020^ cs3941, rabbit anti-pSHP-1^Y564^ D11G5 cs8849, rabbit anti-pErk1/2^T202/Y204^ D13.14.4e cs4370, rabbit anti-pAkt^S473^ 193H12 cs4058, mouse anti-β-Actin 8H10D10 cs3700, mouse anti-LynA+B Lyn01 ab1890, donkey anti-mouse IgG 800CW lc925-32212, donkey anti-rabbit IgG 800CW lc925-32213, donkey anti-mouse IgG 680LT lc925-68022, and donkey anti-goat IgG 680LT lc926-32214. For all data sets, graphing and statistical analyses were performed using Prism software version 10.6.1 (GraphPad, Boston, MA, USA).

### Quantification of RNA and treatment with Actinomycin D

BMDMs were treated, where indicated, with 10 µg/ml Actinomycin D (ActD, Sigma Aldrich) for up to 4 h before lysing in TRIzol (Thermo Fisher, Waltham, MA, USA). RNA was extracted in chloroform (Sigma Aldrich), and purified using RNeasy kits (Qiagen, Hilden, Germany). Synthesis of cDNA was performed using qScript kits (Quantabio, Beverly, MA, USA). SYBR green was used to detect LynA (5’-to-3’ forward (f): ATCCA ACGTC CAA-TAA ACAGCA, reverse (r): ATAAG GCCAC CACAA TGTCAC), LynB (f: TCGAA GACTC AACCA GTTCC TGAA, r: TGAAG GACAA GTCAT CTGGG TG), c-Myc (f: TCCTG TACCT CGTCC GATTC, r: AATTC AGGGA TCTGG TCACG) or Cyclophilin (f: TGCAG GCAAA GACAC CAATG, r: GTGCT CTCCA CCTTC CGT) in 1 µg total RNA via quantitative reverse transcription-polymerase chain reaction (qRT-PCR).

## Results

### Syk phosphorylation in BMDMs is regulated by the total expression of LynA + LynB at steady state

To test whether cumulative Lyn expression or isoform-specificity is the major factor in macrophage signaling, we generated a Lyn-knockout series in which expression levels of LynA and LynB are controlled independently. We previously reported that homozygous single-isoform LynA and LynB knockouts upregulate the remaining isoform of Lyn, LynB or LynA, respectively^10^. These knockouts can therefore be bred to each other for biallelic, supraphysiological expression of the remaining isoform (“UP”) or to complete Lyn^KO^ for monoallelic, WT-like expression of the remaining isoform (“PHYS”) **(Fig.1A)**. We generated this dose series on a (Csk^KO^) Csk^AS^-transgenic background **(Fig.1B)** and tested the ability of each isoform to complement the loss of the other. As on a WT background^10^, homozygous single-isoform knockouts on the Csk^AS^ background upregulated the remaining isoform of Lyn **(Fig.1C)**. Expression of LynA protein in LynA^PHYS^B^KO^ and LynB in LynB^PHYS^A^KO^ protein was comparable to their respective expression levels in WT-Lyn-expressing BMDMs. Although either Lyn isoform could be upregulated when expressed in homozygous knockout mice, LynB **(Fig.1D)** was upregulated almost 2-fold more than LynA **(Fig.1E)**, leading to a higher total LynA + LynB expression in LynB^UP^A^KO^ than in LynA^UP^B^KO^ BMDMs **(Fig.1F)**. Our Lyn-knockout series, therefore, could be used to test whether overexpression of one Lyn isoform could compensate for deletion of the other and evaluate signaling functions across a range of total Lyn expression.

**Figure 1:**
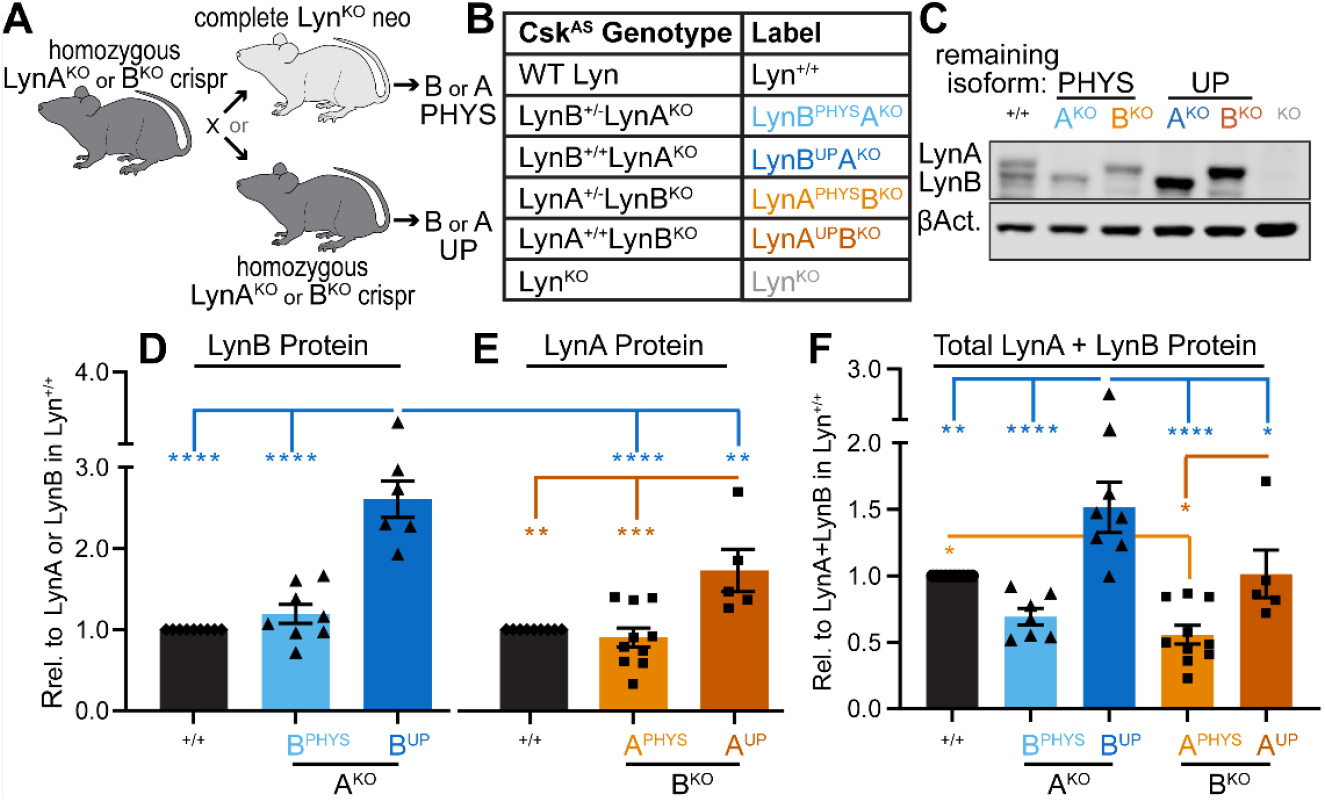
Titration of LynA and LynB expression in BMDMs with a Lyn-knockout genetic series. **(A)** Crossing homozygous single-isoform LynA or LynB knockouts (LynA^KO^ or B^KO^ crispr) to neo-disrupted complete Lyn^KO^ for hemizygous expression or crossing with another homozygous parent for homozygous expression of the remaining isoform, Lyn(B or A), to produce PHYS and UP genotypes, respectively. **(B)** Nomenclature for BMDMs in the Lyn-knockout series; fully LynA- and LynB-expressing Csk^AS^ BMDMs are referred to as Lyn^+/+^. **(C)** Representative immunoblots showing LynA and LynB protein expression in BMDM lysates from the Lyn-knockout series. “Lyn” is omitted from the labels for clarity; β-Actin (βAct.) is shown as a visual loading control.Densitometry quantification of **(D)** LynB protein, **(E)** LynA protein, and **(F)** total LynA + LynB protein in BMDM lysates from the Lyn-knockout series, corrected for the total protein content in each lane and shown relative (Rel.) to levels in Lyn^+/+^.Error bars reflect the standard error of the mean (SEM), with n=11 independent replicates for ^+/+^, 8 for B^PHYS^A^KO^, 7 for B^UP^A^KO^, 10 for A^PHYS^B^KO^, and 5 for A^UP^B^KO^. Significance (Sig.) was assessed using 1-way (1w)-ANOVA with Tukey’s (Tu) test for multiple comparisons: ^****^P≤0.0001, ^***^P=0.0010, ^**^P=0.036-0.0069, ^*^P=0.0112-0.0434. No significant differences (ns) were found for other pairs. In all figures, asterisk color encodes pairwise comparison at a particular time point.

To assess the potential of LynA and LynB to phosphorylate substrates without the complicating factor of the ITAM-organized signalosome, we treated BMDMs from the Lyn-knockout series with a maximally SFK-activating^15^ concentration of the Csk^AS^ inhibitor 3-IB-PP1 **(Fig.2A)**. In agreement with our previous studies^15^, complete knockout of LynA and LynB severely impaired phosphorylation of Syk^Y352^ in response to 3-IB-PP1. We further found that expression of either LynA or LynB alone increased Syk phosphorylation in a dose-dependent manner **(Fig.2B)**.

**Figure 2:**
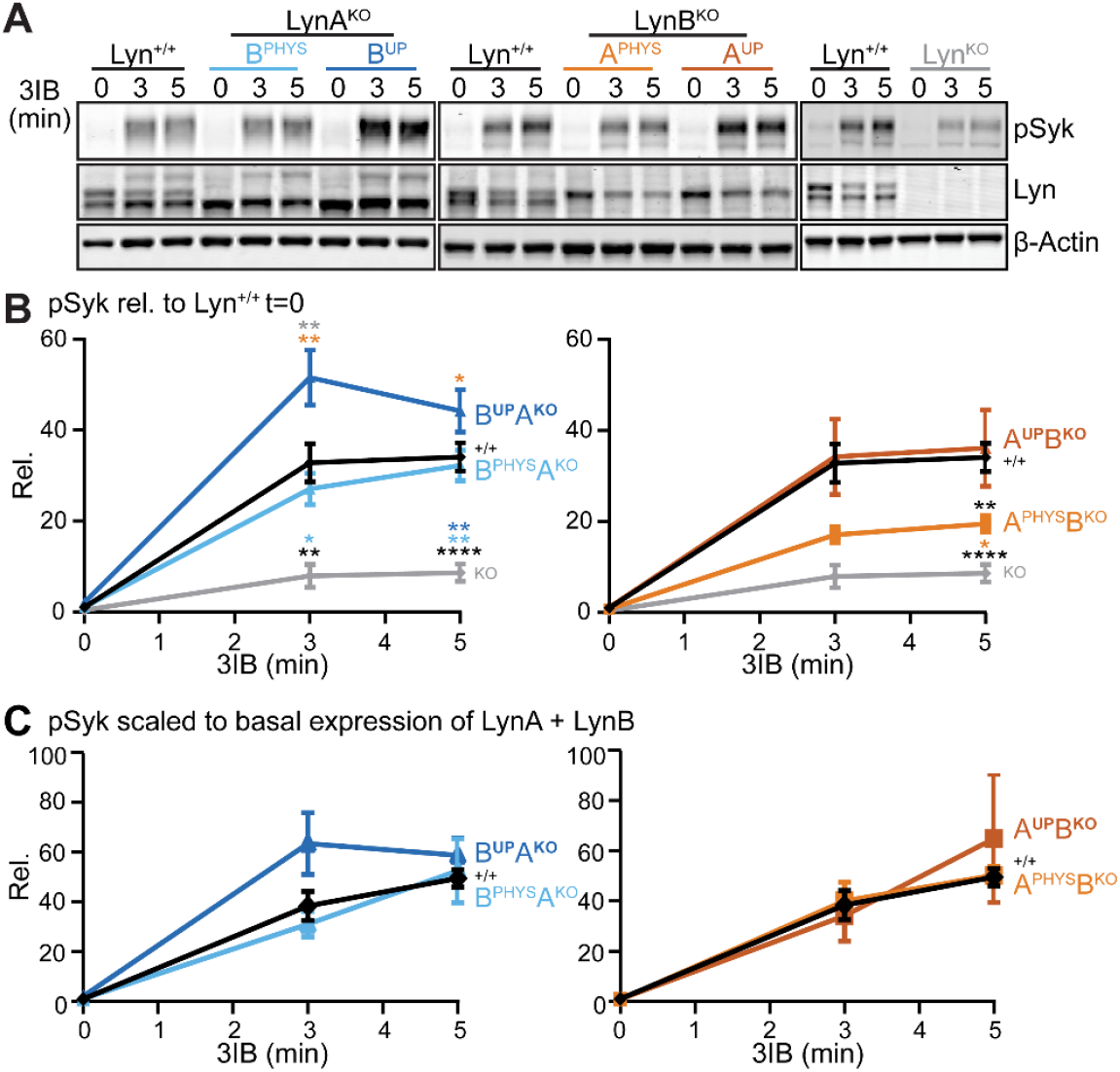
LynA and LynB phosphorylate Syk in a dose-dependent manner: **(A)** Representative immunoblots showing Lyn and pSyk^Y352^ in lysates from Lyn-knockout-series BMDMs treated up to 5 min with 10 µM 3-IB-PP1 (3IB); β-Actin is a visual loading control. Quantification of pSyk^Y352^, corrected for the total protein content in each lane and shown **(B)** relative to levels in Lyn^+/+^ or **(C)** scaled to the basal expression of LynA + LynB. Error bars: SEM, with n=15 for ^+/+^, 9 for A^PHYS^B^KO^, 5 for A^UP^B^KO^, 7 for B^PHYS^A^KO^, 5 for B^UP^A^KO^, and 4 for ^KO^. Sig. assessed using 2-way (2w)-ANOVA-Tu test: ^****^P≤0.0001, ^**^P=0.0017-0.0078, ^*^P=0.0101-0.0329; otherwise ns.

As LynA is degraded 5-fold more rapidly than LynB upon activation^17,18^, we initially hypothesized that differential degradation over time might underlie the lower Syk phosphorylation observed in LynB-knock-out BMDMs. After observing that LynA degradation was unaffected by the absence of LynB **(Fig.S1A)**, we scaled the pSyk data to account for differences in total expression of LynA and LynB over the 3-IB-PP1 treatment time, during which LynA protein would be more quickly degraded than LynB protein. Although the variability of the data eliminated the significant differences in the results, this correction did not cleanly explain the trends in the unscaled data **(Fig.S1B)**.

We then turned to our initial observation that LynB is the more highly expressed isoform at steady state. This is likely a combined effect of steady post-translational loss due to degradation induced gradually by basal phosphorylation of LynA^18^ and regulation at the RNA level **(Fig.S2A)**. As the half-lives of *LynA* and *LynB* mRNA are comparable **(Fig.S2B)**, the relative abundance of *LynB* mRNA is likely due to increased production.

We therefore tested the effects of scaling Syk phosphorylation to the basal expression of LynA + LynB. To our surprise, basal protein levels fully accounted for the signaling amplitude differences between the single-isoform knockouts as well as the dose effects of WT-like or supraphysiological expression of the remaining isoform in the PHYS and UP genotypes **(Fig.2C)**. These data support a model in which LynA and LynB have equal capacity to phosphorylate Syk and the combined basal level of LynA and LynB determines the extent of Syk phosphorylation.

### Dose-dependent effects of Lyn on initiating signaling events are lost downstream of Syk

We next evaluated the effect of Syk phosphorylation on downstream signal propagation, building on our exten-sive recent analysis of 3-IB-PP1-induced signaling in Lyn^KO^ BMDMs^15^. Expression of either LynA or LynB, independently of expression level, was sufficient to suppress basal phosphorylation of Erk and Akt and restore balanced signaling to the Erk and Akt pathways. Interestingly, these downstream effects were quantitatively uncoupled from the extent of upstream Syk phosphorylation **(Fig.3A-B)**. These findings indicate that a modest level of Syk phosphorylation is sufficient to drive downstream pathway activation and that macrophages actively suppress signal propagation once upstream signaling exceeds a defined threshold. A regulatory mechanism, therefore, must prevent excessive pathway output from unrestrained Lyn signaling in the absence of receptor engagement and synapse formation.

**Figure 3:**
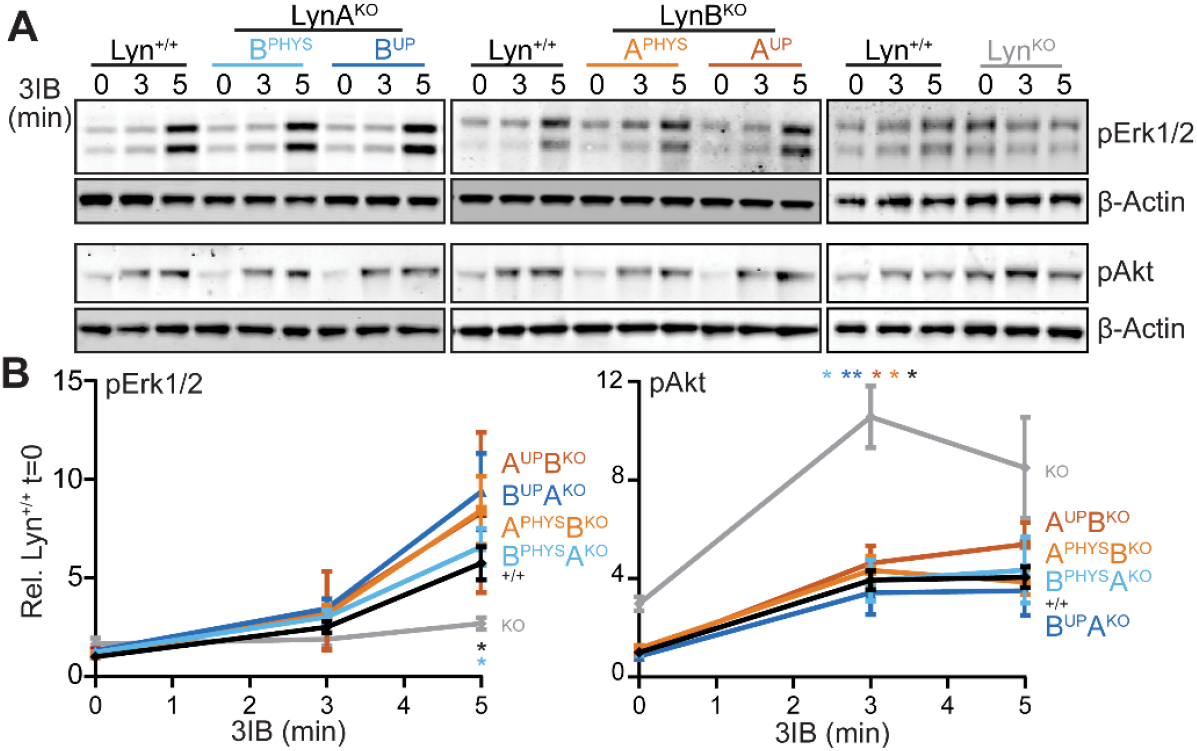
Beyond a threshold of LynA or LynB expression, Erk and Akt signaling are fully restored. **(A)** Representative immunoblots from samples in Figure 2, showing pErk1/2^Y202/T204^ and Akt^S473^; β-Actin is a visual loading control. **(B)** Quantification of pErk1/2^Y202/T204^ and Akt^S473^, corrected for the total protein content in each lane and shown relative to levels in Lyn^+/+^. Error bars: SEM, with n=11 for ^+/+^, 6-8 for B^PHYS^A^KO^, 6-7 for B^UP^A^KO^, 6-9 for A^PHYS^B^KO^, 5-6 for A^UP^B^KO^, and 4-7 for ^KO^. Sig. assessed using 2w-ANOVA-Tu test: ^**^P=0.0087, ^*^P=0.0110-0.0430.

Based on our previous observations that basal Lyn-SHIP1 signaling is a key regulator basal and 3-IB-PP1-induced signaling^15^, we probed SHIP1 phosphorylation in BMDMs from the Lyn-knockout series, at rest and treated with 3-IB-PP1. As with Syk, expression of LynA or LynB restored basal phosphorylation of SHIP1 in a dose-dependent manner **(Fig.4A-B)**. In harmony with our previous study of complete Lyn^KO15^, pharmacological activation of all SFKs induced robust SHIP1 phosphorylation across all genotypes **(Fig.4C)**. This suggests that either isoform of Lyn can phosphorylate SHIP1, and the concomitant increase in basal pSHIP1 counter-acts pSyk to uncouple strong initiating signaling from a downstream antimicrobial response in the absence of receptor engagement.

**Figure 4:**
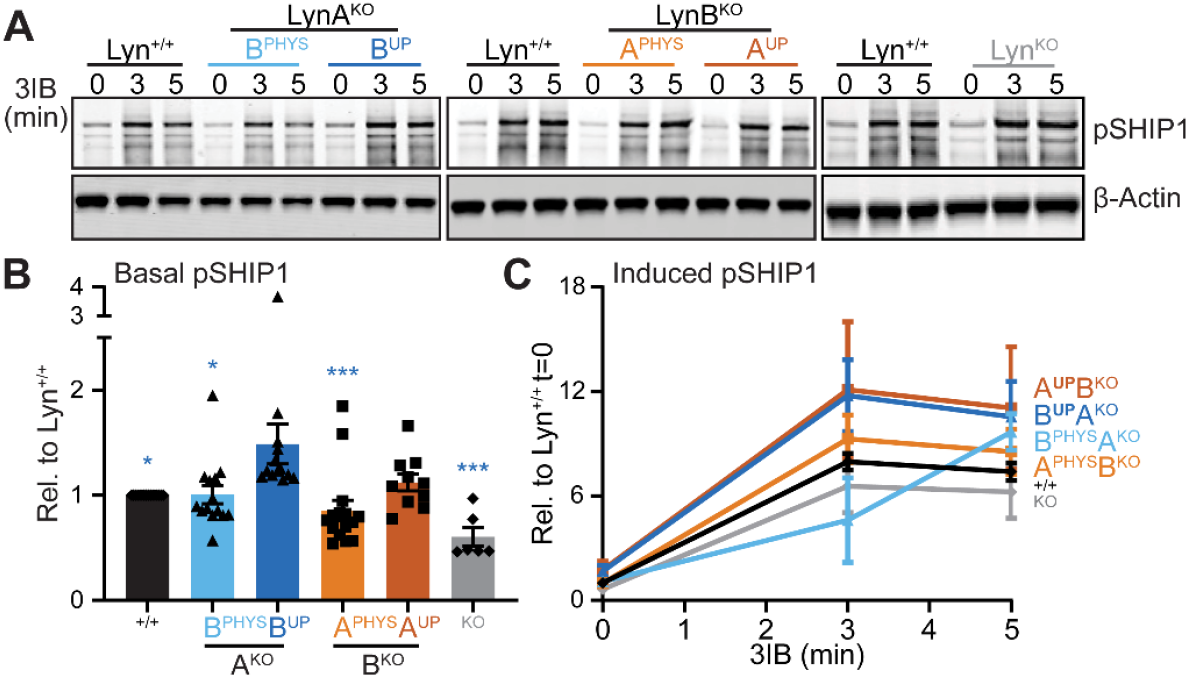
Basal but not induced phosphorylation of SHIP1 is sensitive to the expression levels of LynA and LynB. **(A)** Representative immunoblots from samples in Figure 2, showing pSHIP1^Y1020^; β-Actin is a visual loading control. **(B)** Quantification of basal pSHIP1^Y1020^, corrected for the total protein content in each lane and shown relative to Lyn^+/+^. Error bars in both panels: SEM, with n=11 for ^+/+^, 5-7 for B^PHYS^A^KO^ and B^UP^A^KO^, 5 for A^PHYS^B^KO^ and A^UP^B^KO^, and 4-7 for ^KO^. Sig. assessed using 1w-ANOVA-Tu test: ^***^P=0.0001-0.0006, ^*^P=0.018-0.0182. **(C)** Quantification of pSHIP1 induced by treatment with 3-IB-PP1, corrected for total protein in each lane and shown relative to Lyn^+/+^ t=0. Sig. assessed using 2w-ANOVA-Tu test (ns).

## Discussion

We generated a series of Lyn-knockout mice to uncouple any isoform-specific functions of LynA and LynB from the dose effects of their knockout during pharmacological SFK activation. Our data suggest that both LynA and LynB have the intrinsic ability to initiate ITAM-pathway signaling by phosphorylating Syk and that overexpression of either isoform can compensate for loss of the other. Importantly, our results extend our earlier observation^15^ that basal phosphorylation of SHIP1, largely performed by Lyn, is the key factor that determines the potential activation of downstream signaling. As autophosphorylation site in its unique insert flags LynA for c-Cbl-mediated^17^ polyubiquitination and degradation^18^, we were also able to probe the effects of Lyn degradation on signaling potential—we found that the basal expression of total LynA + LynB better accounts for the difference in degree of Syk phosphorylation than the changes in Lyn protein over time. While this was initially surprising, it does harmonize with the role of Lyn in regulating basal pSHIP1, which both isoforms do in a dose-dependent manner. Therefore, we speculate that LynA degradation could regulate the overall steady-state level of Lyn expression, which in turn could confer cell-type specificity on the sensitivity of different cells to activation. Indeed, we showed previously that mast cells, which express almost no c-Cbl, are highly activatable by 3-IB-PP1, and upregulation of c-Cbl in these cells changes that setpoint^17^; we might now reframe this effect to be one of basal regulation rather than inducible signaling.

We also found that the degree of induced Syk phosphorylation was uncoupled from downstream signal strength. Our previous studies have included an extensive pathway characterization, showing that Lyn expression is necessary for upstream Syk and PLCγ2 phosphorylation, but not PI3K phosphorylation, which promotes balanced Erk and Akt signaling during pharmacological SFK activation^15,18^. It was therefore striking that any level of LynA or LynB expression was sufficient to support WT-like basal regulation of Erk and Akt phosphorylation and induction of both pathways, despite dose-dependent phosphorylation of Syk. This likely reflects the dose-dependent basal phosphorylation of SHIP1, which counters the signal-promoting function of Syk.

Our studies in BMDMs confirm our earlier observation^10^ that macrophages express more LynB than LynA, and we now show that this regulation occurs at the RNA and protein levels. In the Lyn-knockout series, we further observe that homozygous LynB^UP^A^KO^ BMDMs produce higher levels of LynB than LynA^UP^B^KO^ BMDMs do of LynA, again, at the RNA and protein levels. We might have expected the opposite effect because of the design of the CRISPR mutations. LynA-knockout was achieved by deleting part of the LynA unique insert, leading to a premature termination codon and likely nonsense-mediated decay^10^. Therefore, we expected processing of transcript to be split between *LynA* and *LynB* mRNAs, which would not increase *LynB*. In contrast, LynB-knockout was achieved by introducing a point mutation into the exon 2 splice site^10^, a modification designed to bias production of mRNA completely toward *LynA*, which was not enriched as highly as *LynB*. For this study, we need not rely on a particular interpretation to use the varying Lyn expression levels as tools to probe their dose-specific functions. However, it is tempting to speculate that LynB expression is actively regulated to tune the total level of LynA + LynB and therefore basal Lyn and SHIP1 activity. However, it is also formally possible that the CRISPR deletion had an unforeseen effect on the RNA splice site biasing production of *LynB* mRNA. Therefore, we remain cautious about this finding.Regardless of the origin, our study suggests that LynA and LynB have complementary functions and that the increased severity of autoimmunity in LynB-knockout (LynA^PHYS^B^KO^) mice^10^ may have been due to the lower overall expression of LynA + LynB protein in this genotype compared to LynB^PHYS^A^KO^.

While this study focused on the intrinsic potential of LynA and LynB to phosphorylate their substrate Syk and initiate signaling, isoform-specific functions may be revealed in other signaling pathways or contexts. Although we have also found that LynA and LynB can both suppress macrophage responses to TLR4 and TLR7^25^, other pathways may yet be regulated differentially by the two Lyn isoforms.

In summary, our findings support a model in which total LynA + LynB expression effects a basal inhibitory program by phosphorylating SHIP1, restricting macrophage activation by activated SFKs in the absence of particulate receptor engagement and a phagocytic synapse. Increased inhibitory signaling in the macrophage steady state creates a bottleneck between enhanced initiating events and productive signal propagation.

## Data availability

The data that support the findings of this study are available from the corresponding author upon request.

## Authorship Roles

Conceptualization-Equal (SES, TSF), Data Curation-Equal (SES, TSF), Formal analysis-Equal (SES, TSF), Funding acquisition-Lead (TSF), Funding acquisition-Supporting (CES), Investigation-Lead (SES), Investigation-Supporting (CES, STR), Methodology-Equal (SES, TSF), Methodology-Supporting (CES), Project administration-Lead (TSF), Resources-Lead (TSF), Supervision-Lead (TSF), Supervision-Supporting (SES), Validation-Lead (TSF), Visualization-Equal (SES, TSF), Writing-original draft-Equal (SES, TSF), Writing-reviewing and editing-Equal (SES, TSF), Writing-reviewing and editing-Supporting (CES, STR).

## Acknowledgements

We thank Drs. Kaylee Schwertfeger and Jennifer Tuokkola for qRT-PCR training and instrument use. Many thanks also to Drs. Anders Lindstedt and William Kanagy for valuable feedback and discussion and to Dr. Ben Brian for the mouse cartoon. Thanks to Monica Sauer for mentoring junior lab members and to Dr. J.T. Greene for helping to design the LynB qRT-PCR primers.

## Figure Legends

**Figure S1: Activation-induced degradation of LynA protein does not explain differences between LynA and LynB function in Syk phosphorylation. (A)** LynB and LynA protein levels over time in lysates from BMDMs in the Lyn-knockout series treated with 10 µM 3-IB-PP1. **(B)** Quantification of pSyk^Y352^ scaled to the expression of LynA + LynB over time. In both panels, error bars: SEM, with n=11 for ^+/+^, 6 for A^KO^ genotypes, and 9 for B^KO^ genotypes. Sig. assessed using 2w-ANOVA-Tu test (ns).

**Figure S2: Differences in RNA production contribute to differences in LynA and LynB protein expression in BMDMs. (A)** qRT-PCR quantification of *LynB* and *LynA* mRNA in BMDM lysates from the Lyn-knockout series, corrected for *cyclophilin (Ppia)* expression and shown relative to expression in WT. *LynA* and *LynB* mRNAs were undetectable in their respective knockouts. Error bars: SEM, with n=11-13 for WT and UP genotypes and 3 for PHYS genotypes. Sig. assessed using 1w-ANOVA-Tu test: ^****^P≤0.0001, ^**^P=0.0016-0.0071, ^*^P=0.0120-0.0400. **(B)** mRNA degradation in WT BMDMs during ActD treatment, fit to a single-exponential decay; rapidly-degrading c-Myc is a control for ActD efficacy. Error bars: SEM, n=3. Sig. assessed using 2w-ANOVA-Tu test: ^***^P=0.0005, ^**^P=0.0014-0.0093, ^*^P=0.0217.

**Figure S1:**
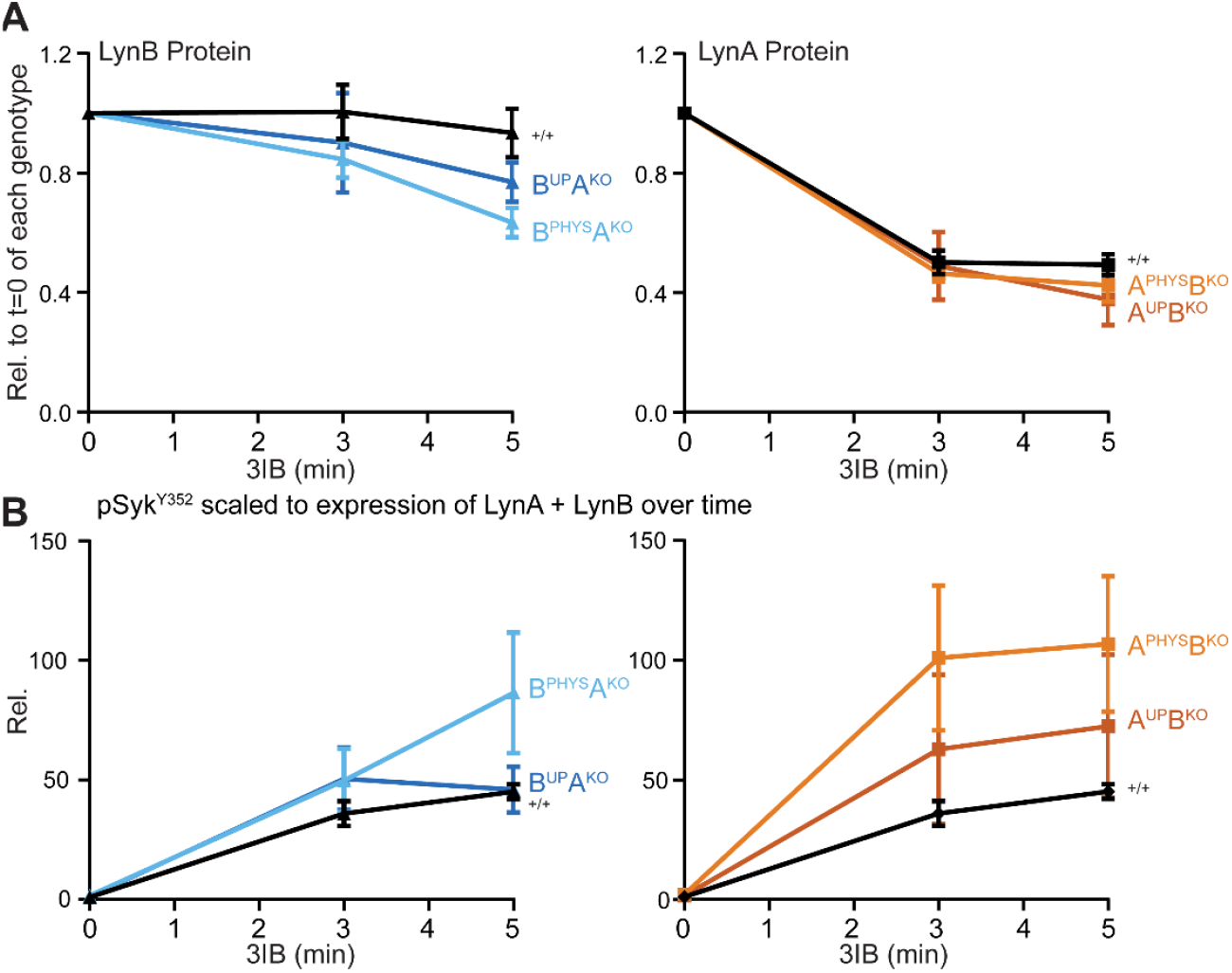
Activation-induced degradation of LynA protein does not explain differences between LynA and LynB function in Syk phosphorylation. **(A)** LynB and LynA protein levels over time in lysates from BMDMs in the Lyn-knockout series treated with 10 µM 3-IB-PP1. **(B)** Quantification of pSyk^Y352^ scaled to the expression of LynA + LynB over time. In both panels, error bars: SEM, with n=11 for ^+/+^, 6 for A^KO^ genotypes, and 9 for B^KO^ genotypes. Sig. assessed using 2w-ANOVA-Tu test (ns within any of the four graphed date sets).

**Figure S2:**
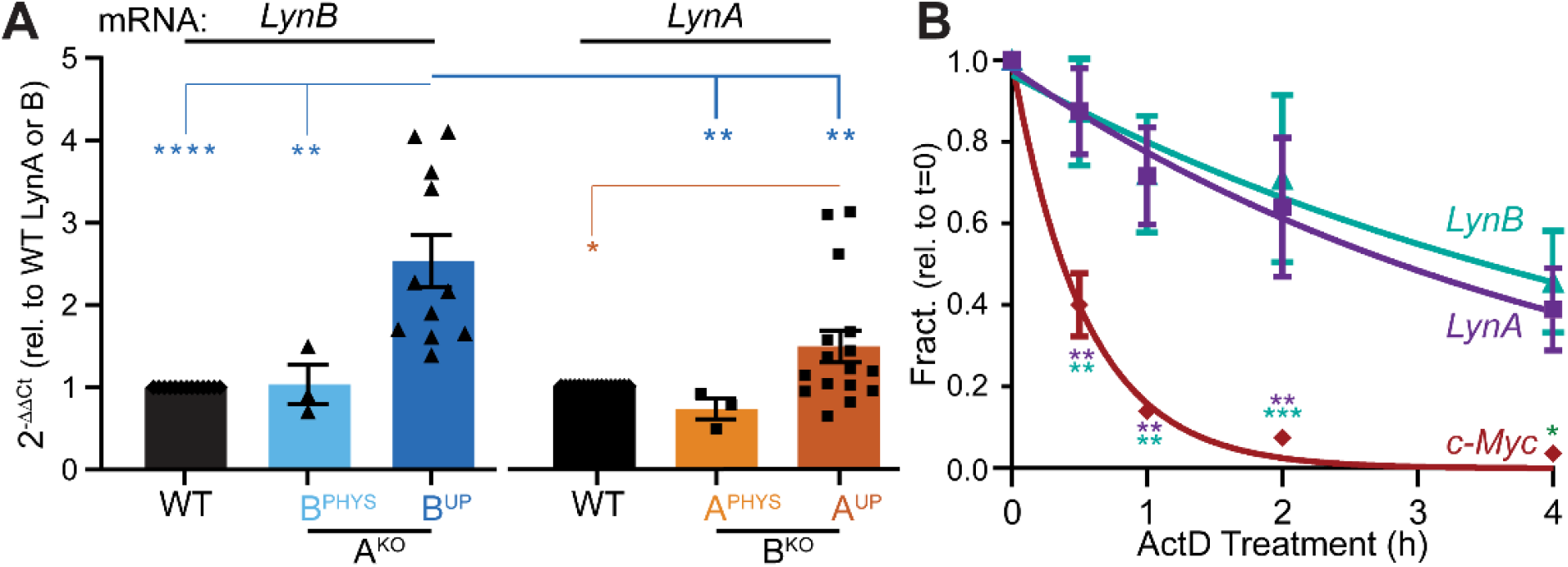
Differences in RNA production contribute to differences in LynA and LynB protein expression in BMDMs. **(A)** qRT-PCR quantification of *LynB* and *LynA* mRNA in BMDM lysates from the Lyn-knockout series, corrected for *cyclophilin (Ppia)* expression and shown relative to expression in WT. *LynA* and *LynB* mRNAs were undetectable in their respective knockouts. Error bars: SEM, with n=11-13 for WT and UP genotypes and 3 for PHYS genotypes. Sig. assessed using 1w-ANOVA-Tu test: ^****^P≤0.0001, ^***^P=0.0010 ^**^P=0.016-0.0053, ^*^P=0.0120-0.0400. **(B)** mRNA degradation in WT BMDMs during ActD treatment, fit to a single-exponential decay; rapidly-degrading *c-Myc* is a control for ActD efficacy. Error bars: SEM, n=3. Sig. assessed using 2w-ANOVA-Tu test: ^***^P<0.001, ^**^P<0.01, ^*^P<0.05.

